# Human gaze behaviors track abstract stimulus categories

**DOI:** 10.1101/2025.08.08.669367

**Authors:** Ali Caron, Edward F. Ester

## Abstract

Categorization, or the ability to group stimuli according to behavioral relevance, is a cornerstone of abstract cognition. Neurophysiological studies in non-human primates have revealed that category-selective signals are robustly encoded in oculomotor structures, including the lateral intraparietal area (LIP) and superior colliculus (SC), and that this encoding produces small, uninstructed eye movements that reflect learned category distinctions. Whether a similar phenomenon exists in humans is unknown. Here, we show that human gaze behavior encodes abstract categorical information independent of physical stimulus features. In three experiments, participants learned to classify oriented stimuli according to an arbitrary rule while their eye movements were recorded. Category identity could be reliably decoded from records of gaze position, particularly on trials with accurate categorization. A delayed match-to-category task confirmed that category-selective gaze patterns emerged before any motor response could be planned, ruling out response-related confounds. A further experiment demonstrated that gaze patterns tracked both orientation and color categories, confirming that the decoded signal reflects the observer’s behavioral state rather than stimulus-evoked oculomotor biases. These findings establish that human oculomotor behavior carries abstract cognitive signals, paralleling recent non-human primate results linking incidental gaze shifts to the multiplexed encoding of category information within oculomotor networks.

## Introduction

Categorization describes the ability to group stimuli, events, and actions according to their behavioral relevance (Ashby & Maddox, 2005). Laboratory studies of rule-based categorization have revealed robust category-selective signals across a distributed network of cortical and subcortical regions (e.g., Freedman et al., 2001; Freedman & Assad, 2006; Swaminathan & Freedman, 2012; Ester et al., 2020; Peysakhovich et al., 2024; Henderson et al., 2025). In non-human primates, neurons encoding category membership have been identified not only in association cortices but also in structures traditionally associated with sensorimotor control, challenging strictly modular views of cognitive function and suggesting that abstract decision variables are embedded within circuits that also control overt behavior.

Recent work in oculomotor systems provides a striking example. The superior colliculus (SC)—a midbrain structure classically implicated in saccade generation—exhibits robust, short-latency encoding of abstract visual categories that can exceed that observed in posterior parietal cortex, and reversible inactivation of SC selectively impairs categorical judgments, implying a causal contribution to abstract decision making (Peysakhovich et al., 2024). Rosen and Freedman (2025a) showed that SC neurons carry multiplexed cognitive and saccadic signals: population activity related to category decisions transiently aligns with saccade-coding subspaces, and this alignment is accompanied by small, uninstructed eye movements that reflect learned category boundaries.

Converging evidence in humans indicates that the oculomotor system is likewise sensitive to internal cognitive states. During visual working memory (WM) tasks, participants make small but systematic eye movements toward the remembered locations of stimuli, even when spatial position is task-irrelevant (van Ede et al., 2019; Ester & Weese, 2023). Fixational eye movements can also carry information about non-spatial properties: Linde-Domingo and Spitzer (2024) demonstrated that miniature gaze patterns encode the orientations of stimuli held in WM, and that their representational geometry evolves from stimulus-specific to more abstract formats across encoding and maintenance. Together, these studies establish that human gaze dynamics provide a sensitive readout of internal sensory representations (see also de Vries & van Ede, 2024; Chota et al., 2025).

Despite these advances, it remains unknown whether human gaze behavior also reflects the abstract, rule-defined categorical status of stimuli, rather than only their physical or remembered features. The oculomotor system appears sensitive to the contents of internal visual representations, but it is not yet clear whether observers’ categorical decision state—defined relative to an arbitrary boundary in feature space—can be recovered from eye movements alone. Addressing this question would provide a behavioral bridge between non-human primate neurophysiology, which links multiplexed category codes in SC to incidental eye movements (Rosen & Freedman, 2025a), and human cognition, while testing the extent to which abstract decision variables are embedded within human oculomotor behavior (Rosen & Freedman, 2025b; Kiyonaga & Serences, 2025).

Here we test whether human oculomotor behavior carries abstract, rule-defined category information independent of low-level stimulus properties and response preparation. Across three experiments, participants classified oriented stimuli into discrete groups defined by an arbitrary, participant-specific boundary while their eye movements were recorded. In Experiment 1, we asked whether stimulus category could be decoded from gaze position during a speeded categorization task. In Experiment 2, we used a delayed match-to-category design to dissociate category encoding from motor preparation. In Experiment 3, we presented identical displays in two tasks—orientation categorization and a color majority judgment—to ask whether gaze-based decoding reflects task-dependent category representations rather than stimulus-driven oculomotor biases. Together, these experiments demonstrate that human gaze dynamics carry a reliable signature of the observer’s categorical decision state, paralleling findings in non-human primates and supporting the view that abstract cognitive variables are multiplexed within oculomotor control circuits.

## Methods

### General Information Common to All Experiments

#### Participants

Eighty volunteers (both sexes; aged 18–40 years; normal or corrected-to-normal visual acuity) participated in one of three experiments (N = 34, 27, 16 for Experiments 1–3, respectively) for course credit or $15/h remuneration. Participants were recruited from the University of Nevada, Reno community. We did not collect demographic data, as we had no a priori reason to expect effects of gender, race, ethnicity, or similar characteristics. Sample sizes were not determined by formal power analyses, as no prior work existed to estimate the expected effect size for category decoding from gaze position; instead, we targeted sample sizes commonly used in within-participant decoding studies of eye movements and working memory (e.g., van Ede et al., 2019).

##### Testing Environment and Stimuli

Stimuli were generated in MATLAB using Psychtoolbox (Brainard, 1997) and presented on a 27-inch G-Sync LCD monitor (240 Hz refresh rate; 1920 × 1080 resolution). Participants sat 85 cm from the display with head position stabilized by an SR Research chinrest (1° visual angle ≈ 49 pixels).

Stimuli were circular apertures (2–9° radius) containing 350 iso-oriented bars (1° × 6 pixels stroke width), flickering at 30 Hz (50% duty cycle) with random replotting per cycle to prevent foveation of individual bars. Category judgments were made via keyboard (“Z” = Category 1; “?” = Category 2).

##### Training Task

For all three experiments, volunteers learned to classify orientation stimuli into discrete groups according to an arbitrary (i.e., experimenter-chosen) boundary by completing a training task. Volunteers viewed apertures of iso-oriented bars and reported whether they were members of “Category 1” or “Category 2” by pressing the “Z” or “?” key (respectively) at any point during each 3 second trial (Figure S1A). Category membership was defined by a boundary whose orientation was randomly and independently selected for each volunteer on the interval [0°, 179°). We defined a set of stimuli tilted ±15°, ±30°, ±45°, and ±60° relative to each volunteer’s category boundary and assigned stimuli tilted counterclockwise to the boundary to Category 1 and stimuli tilted clockwise to the boundary to Category 2 (Figure S1B and 1A). This individualized training ensured that category labels could not be inferred from any fixed retinal orientation but instead had to be learned relative to a participant-specific decision boundary. Feedback – i.e., “correct!” vs. “incorrect!” was given at the end of each trial. Failures to respond within the 3-second stimulus window were treated as incorrect. Volunteers received verbal instructions describing key features of the task. They were told that there existed an (unknown) boundary that divided stimuli into different categories, and that their task was to identify this boundary through trial-and-error. In general, with the benefit of these instructions, volunteers reached asymptotic performance by the 2^nd^ or 3^rd^ block, e.g., Figure S1C-E). Each volunteer completed 3 blocks of 32 trials in this task. Volunteers then completed one of the three experiments described below using the same (bespoke) category boundary imposed during training. Due to a coding error, volunteers in Experiment 2 were trained using stimuli tilted ±15°, ±30°, ±45°, ±60°, and ±75° from the category boundary (instead of ±15°, ±30°, ±45°, and ±60°), but all other aspects of the task were identical to the training regimens used in Experiments 1 and 3.

##### Eyetracker Acquisition and Preprocessing

Binocular gaze position data were recorded via an SR Research Eyelink 1000 Plus infrared eyetracking system. Eyetracker recordings were obtained after applying a 9-point calibration procedure, and a drift correction procedure was applied at the beginning of each experimental block. Re-calibration was performed when necessary, for example, if the volunteer left the testing environment to use the restroom. Recordings were digitized at 500 Hz or 1 kHz.

Gaze data were minimally preprocessed. Data were downsampled from 1 kHz to 500 Hz when necessary and filtered for blinks. Blinks were defined as periods where the recorded pupil size fell below the median pupil size recorded across all experimental trials and conditions. We additionally discarded data from 50 ms before and 50 ms after each blink. Missing data were linearly interpolated, and the resulting gaze timeseries were smoothed with a gaussian kernel (10 ms full-width half-maximum). Saccades were identified using a velocity-based thresholding algorithm described in other work (e.g., Liu et al., 2024). Briefly, gaze position recordings were transformed into a velocity time course by calculating the Euclidean difference between successive samples. Saccades were identified as periods when gaze velocity exceeded 150°/s. All findings reported here generalized across saccade thresholds ranging from 30° to 300°/s.

##### Classification of Categories from Gaze Data

Our primary analysis asked whether trial-wise gaze position carried sufficient information to distinguish between stimulus categories. We used a decoding approach to determine the extent to which records of volunteers [x,y] gaze position differentiated between stimulus categories.

For time-resolved decoding, we labeled trial-wise records of volunteers’ gaze position according to category membership (i.e., Category 1 vs. Category 2). We implemented a leave-one-block-out cross-validation procedure, where gaze position records from N-1 blocks were used to train a linear discriminant classifier (via MATLAB’s “classify” function) and the held-out block was used for testing. We used a linear discriminant function to minimize computational overhead; since decoding was performed over time, training a support vector machine-based classifier took ∼24 hours to complete for one subject, even with 32x parallelization. This procedure was repeated until all experimental blocks had been used as the test data set, and results were averaged across permutations. Decoding was performed on a timepoint-by-timepoint basis using minimally preprocessed data (see above).

For epoch-based decoding, we first extracted fixations using the velocity-based threshold procedure described in the preceding section. Fixations were labeled according to category membership and used to construct training and test data sets containing 80% and 20% of all fixations within an epoch of interest (e.g., sample period, delay period, etc.), respectively. The training data set was used to condition a support vector machine classifier (MATLAB’s “fitcsvm” function), and the trained classifier was used to predict a category label for each observation in the test data set (MATLAB’s “predict” function). We repeated this procedure 20 times while choosing different (randomly selected) subsets of data to define the training and test data sets and averaged the results over permutations. Epoch-based decoding was computed separately using fixations recorded within the outer perimeter of the stimulus aperture (9 DVA) or fixations recorded inside the inner perimeter of the stimulus aperture (i.e., microsaccades; 2 DVA). Generally, this had little effect on decoding accuracy.

During Experiment 1, we omitted ambiguous trials (i.e., those where the presented orientation matched the volunteer’s category boundary) from category decoding analyses. Data from these trials, however, were used to test the possibility that successful category decoding could be related to response demands (for example, looking at the keyboard key or response hand needed to produce the appropriate response on a given trial). To test that possibility, we trained a linear discriminant classifier to distinguish between categories based on records of gaze position measured during unambiguous trials, then applied the trained classifier to records of volunteers choice behavior (i.e., Category 1 or Category 2) during ambiguous trials. We reasoned that if decoding accuracy was driven by response selection or motor activity, then the trained classifier should be able to predict volunteers’ responses during ambiguous trials when no category information was available.

##### Distance-to-Boundary Decoding Analysis

To test whether category decoding reflected a continuous gradient of stimulus orientation or discrete category representations, we performed a distance-to-boundary analysis using data from Experiment 1 (Figure 2E). Stimuli were binned according to their angular distance from each participant’s category boundary (2°, 5°, 15°, or 45°). A linear support vector machine classifier was trained to distinguish Category 1 from Category 2 using saccade endpoints measured at any point during unambiguous trials (i.e., excluding 0° boundary trials), following the leave-one-block-out cross-validation procedure described above. The trained classifier was then applied separately to gaze data from each distance bin to quantify decoding accuracy as a function of angular distance from the boundary. Statistical significance was assessed using bootstrap tests (10,000 permutations).

**Figure 1.**
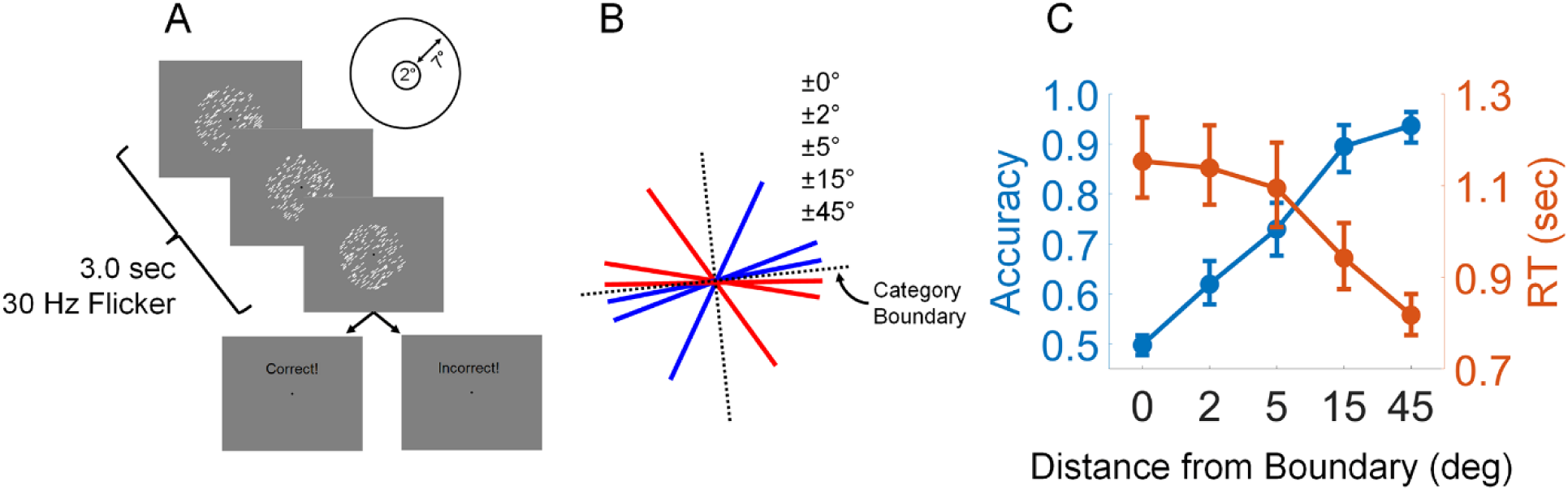
Overview of Experiment 1. (A) Volunteers viewed displays containing circular apertures of iso-oriented bars. (B) Volunteers classified stimulus orientations into discrete groups based on an arbitrary (experimenter-imposed) category boundary. Volunteers could respond at any point during the 3 second stimulus window, and feedback (correct, incorrect) was given at the end of each trial. Note that this schematic is for illustrative purposes and does not contain all possible stimulus orientations (i.e., ±0° and ±2°) (C) Volunteers’ task performance scaled inversely with the angular difference between presented orientations and the category-defining boundary. Error bars depict 95% confidence intervals of the mean.

**Figure 2.**
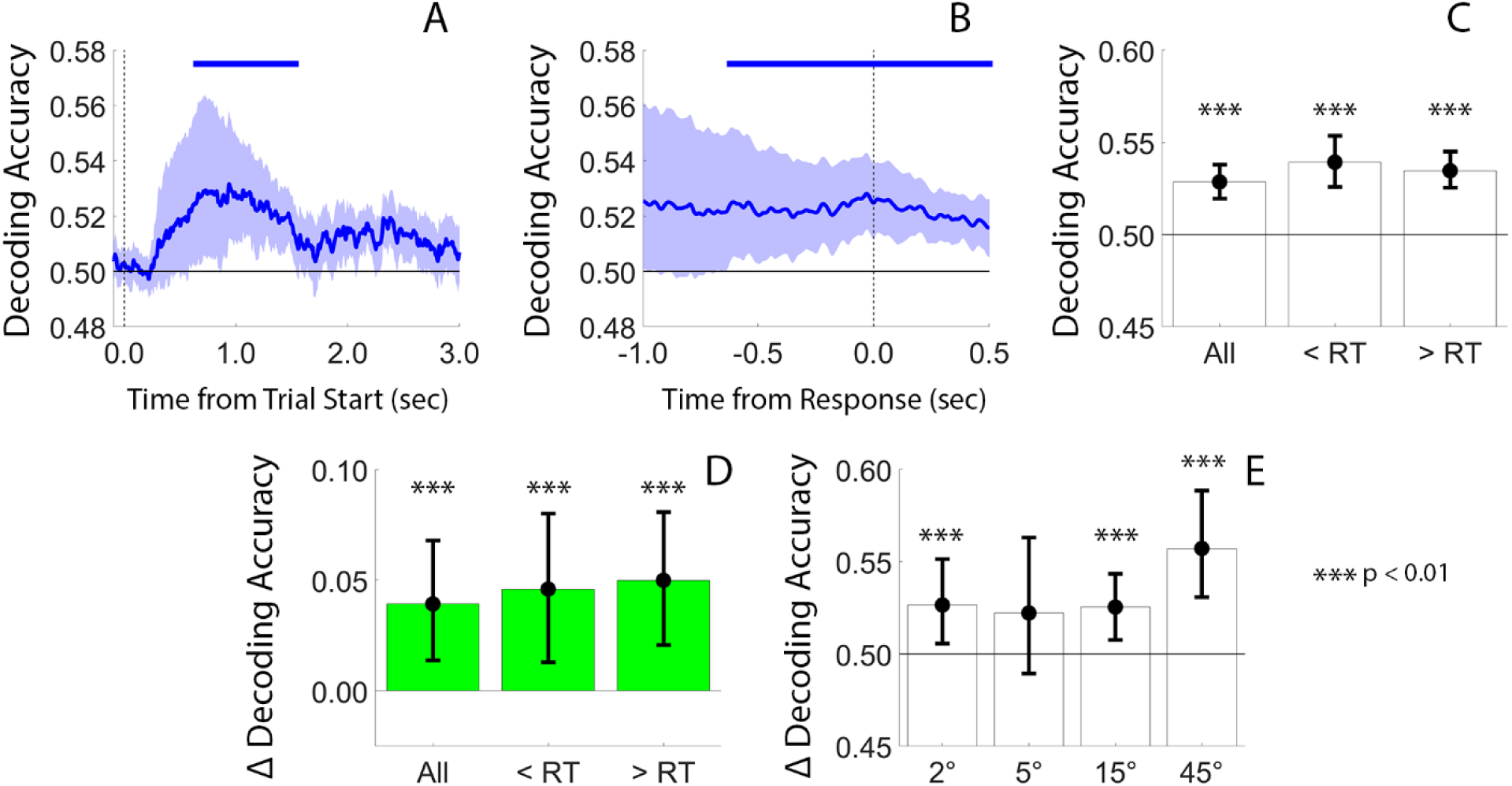
Successful Decoding of Category Membership from Gaze Position. (A) We trained a linear discriminant classifier to predict stimulus category (i.e., Category 1 or Category 2) from volunteers gaze data. Decoding accuracy was modestly but robustly above chance during a period spanning ∼0.8-1.1 sec after trial start, when most responses occurred (see Figure 1C). (B) Category decoding accuracy remained constant over a period spanning -1.0 to +0.5 sec around volunteers’ responses. (C) Category decoding accuracy calculated from fixations during different trial epochs. “All” refers to fixations measured at any point during the 3.0 second stimulus interval, “< RT” refers to fixations measured at any point before a response time threshold, defined as the 80^th^ percentile of each volunteer’s response time distribution calculated over all trials and conditions), and “> RT” refers to fixations measured at any point after the response time threshold. (D) Differences in category decoding accuracy for classifiers trained on fixations measured during correct trials vs. fixations measured during all trials, i.e., correct trials minus all trials. (E) Category decoding performance sorted by the absolute angular distance between the presented stimulus and each participant’s category boundary. Shaded areas and error bars in each panel depict the 95% confidence interval of the mean. The horizontal lines at 0.5 depict chance decoding accuracy. The horizontal bars at the top of panels A and B depict epochs where category decoding accuracy exceeded chance levels (nonparametric signed randomization tests; Methods).

##### Trial-wise Logistic Regression

To determine which combinations of eye-movement parameters predicted trial-wise category membership, we fit a mixed-effects logistic regression model with stimulus category (Category 1 vs. Category 2) as the binary dependent variable. Fixed-effect predictors were: (1) total saccade count per trial, (2) mean saccade amplitude per trial, (3) mean vertical saccade endpoint coordinate per trial (sine-transformed), and (4) mean horizontal saccade endpoint coordinate per trial (cosine-transformed). Participant was included as a random intercept. Saccades were identified using the velocity-threshold procedure described above. Fixed-effect coefficients were tested for significance using t-tests, with false-discovery rate correction for multiple comparisons across the four predictors. This analysis was performed using data from Experiment 1 (Table 1) and separately for Experiment 2 (Table 2).

**Table 1.**
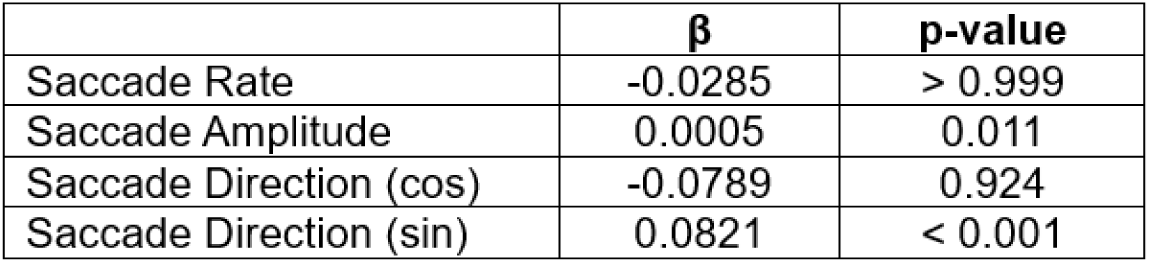
Results of a mixed-effects logistic regression linking eye-movement parameters (saccade rate, mean amplitude, and mean horizontal and vertical direction) to stimulus category in Experiment 1.

| | $\beta$ | p-value |
| --- | --- | --- |
| Saccade Rate | -0.0285 | > 0.999 |
| Saccade Amplitude | 0.0005 | 0.011 |
| Saccade Direction (cos) | -0.0789 | 0.924 |
| Saccade Direction (sin) | 0.0821 | < 0.001 |

**Table 2.** Results of a mixed-effects logistic regression linking eye-movement parameters (saccade rate, mean amplitude, and mean horizontal and vertical direction) to sample category in the delayed match-to-category task (Experiment 2).

| | $\beta$ | p-value |
| --- | --- | --- |
| Saccade Rate | 4.657 | < 0.001 |
| Saccade Amplitude | -0.002 | 0.999 |
| Saccade Direction (cos) | -0.134 | 0.999 |
| Saccade Direction (sin) | -0.051 | 0.695 |

##### Color Decoding Analyses

Finally, in Experiment 3 we tested whether gaze patterns also carried information about a non-spatial, non-orientation feature dimension (color), and whether this signal depended on the current task. We trained support vector machine classifiers (fitcsvm, MATLAB) to decode “color category” from fixation locations during Experiment 3 (Figure 5F-G). Color category was defined by the proportion of white bars in each display. Displays containing >50% white bars were assigned to one color category, and displays containing >50% black bars were assigned to the other, yielding balanced categories. Fixations were extracted using the velocity-threshold procedure and labeled according to color category. We used an 80/20 train/test split with 20-fold permutation testing, separately for data from the orientation categorization task and color categorization task. Fixations falling outside the 9° stimulus aperture were excluded. Population-level decoding accuracy was assessed using bootstrap tests (10,000 resamples), with task differences evaluated by comparing the proportion of bootstrap samples where decoding accuracy differed between tasks.

##### Statistical Tests

Statistical analyses of time-resolved decoding accuracy were based on nonparametric signed randomization tests (Maris & Oostenveld, 2007). Unless otherwise specified, each test we performed assumes a null statistic of 50%. We therefore generated null distributions by randomly relabeling each volunteer’s data with 50% probability and averaging the data across volunteers. This step was repeated 10,000 times, yielding a 10,000-element null distribution for each time point. Finally, we implemented a cluster-based permutation test with cluster-forming and cluster size thresholds of *p* < .05 (two-tailed) to evaluate observed differences with respect to the null distribution while accounting for signal autocorrelation.

Statistical analyses of epoch-based decoding accuracy were based on bootstrap tests. We simulated population-level decoding accuracy by randomly selecting (with replacement) and averaging data from N of N volunteers (where N is the total number of volunteers in each experiment) 10,000 times. We then calculated the proportion of permutations where each sample mean was less than or equal to chance decoding accuracy (50%) or 0 (e.g., for analyses comparing decoding accuracy during correct vs. all trials). We also used these distributions to calculate 95% confidence intervals for each statistic; these are the error bars shown in the figures below.

We quantified category-level differences in the distributions of volunteers’ gaze position (e.g., Figure 3D) using two-sample Kuiper tests (i.e., the circular analog of the Kolmogorov-Smirnov test, which evaluates whether two samples are drawn from the same underlying distribution). Volunteer and category-specific gaze coordinates were converted to polar format and projected onto a unit circle. Because each volunteer was assigned a unique category boundary, we calculated the Kuiper statistic separately for each volunteer and recorded the proportion of positive outcomes, indicating that a volunteer’s fixation coordinates are significantly different during Category 1 vs. Category 2 trials.

**Figure 3.**
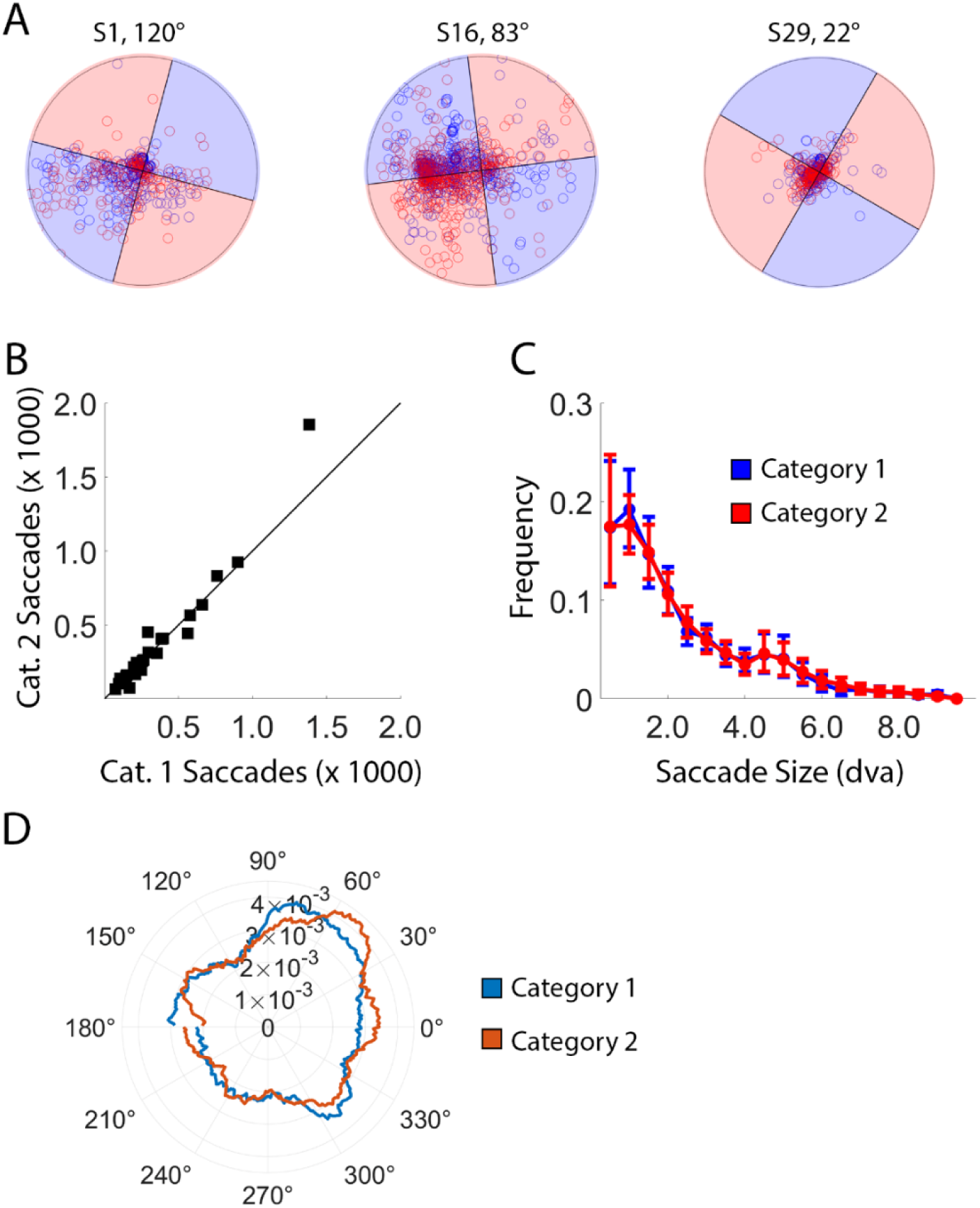
Oculomotor Behaviors During Experiment 1. (A) Plots of individual fixations recorded across the entire experimental testing session for three representative volunteers with category boundaries of 120°, 83°, and 22°, respectively. Blue and red circles reflect fixations during Category 1 and Category 2 trials, while the shaded areas in each plot depict the range of orientations assigned to Categories 1 and 2 for each volunteer. The outer radius of each plot corresponds to the outer radius of the stimulus aperture, i.e., 9 DVA. (B) Total fixation counts during Category 1 and Category 2 trials, plotted against a unity line. Each square is an individual volunteer. (C) Histogram of fixation amplitudes. Most fixations were small, within ∼2-3° of the display center. (D) Polar angles of fixations plotted as a function of category membership. Eye position data from each volunteer were aligned to a common boundary of 0°; thus, in this plot values from 0° to 90° and 180° to 270 degrees correspond to Category 1 orientations, while values from 90° to 180° and 270° to 359° correspond to Category 2 orientations.

##### Data & Code Availability

Raw data, stimulus presentation software, and analytic software sufficient to produce each manuscript figure will be made publicly available upon publication of this paper.

### Experiment-Specific Information

#### Experiment 1

##### Data Exclusion

Data from 6 of the 34 volunteers who completed Experiment 1 were discarded. One volunteer withdrew from the study after completing only 2 experimental blocks. Data from 5 other volunteers were discarded due to hardware or software faults (e.g., codes marking trial events of interest were not sent from the stimulus presentation computer to the eyetracker recording computer, or monocular instead of binocular recordings were obtained). Thus, the data reported here reflect the remaining 28 volunteers.

##### Experimental Task

A trial schematic is shown in Figure 1A-B. We defined a set of oriented stimuli ±2°, ±5°, ±15°, and ±45° relative to the volunteer’s category boundary, with negative values assigned to Category 1 and positive values assigned to Category 2. Volunteers reported the category membership of presented stimuli by pressing the “z” and “?” keys (for Categories 1 and 2, respectively) at any point during each three-second trial. Failures to respond within the 3 second trial window were treated as incorrect. We also included trials where the presented stimulus precisely matched the category boundary, i.e., ±0°. Volunteers were not informed of this manipulation. To maintain the ruse, volunteers were told to guess if they were uncertain, and random feedback (i.e., correct vs. incorrect) was given at the end of these trials. Volunteers completed 9 (N = 1), 11 (N = 1), 16 (N = 1), 17 (N = 1), and 18 (N = 24) blocks of 36 trials.

#### Experiment 2

*Data Exclusion*. Data from 2 of the 27 volunteers who completed Experiment 2 were discarded. One volunteer withdrew from the study after completing only 1 experimental block, and data from a second volunteer were discarded due to a hardware fault that occurred midway through testing. Thus, the data reported here reflect the remaining 25 volunteers.

##### Experimental Task

A task schematic is shown in Figure 4A-B. Volunteers viewed successive displays of sample and test stimuli (500 ms each) separated by a blank interval (1000 ms). We defined a set of oriented stimuli ±15°, ±30°, ±45°, ±60° and ±75° relative to the volunteer’s category boundary, with negative values assigned to Category 1 and positive values assigned to Category 2. Volunteers were required to report whether the sample and test stimuli were drawn from the same category, pressing “z” to indicate “yes” and “?” to indicate “no”. Volunteers could respond at any point during a 3 second window beginning with the test display, and failures to respond within this window were treated as incorrect. The sample and test stimuli could be drawn from either category, with the exception that the same stimulus value (e.g., +15°) could not be presented as both the sample and the test. This task decouples categorization from response preparation across clearly defined temporal epochs: the sample period (0–500 ms), the delay period (500–1500 ms), and the test period (1500–4500 ms). Since volunteers cannot know whether the sample and test belong to the same category until the test appears, any category-selective signals measured during the sample and delay periods (0–1500 ms) cannot be ascribed to response selection or motor preparation. Volunteers completed 9 (N = 2), 13 (N = 1), 14 (N = 1), 17 (N = 1) and 18 (N = 20) blocks of 40 trials.

**Figure 4.**
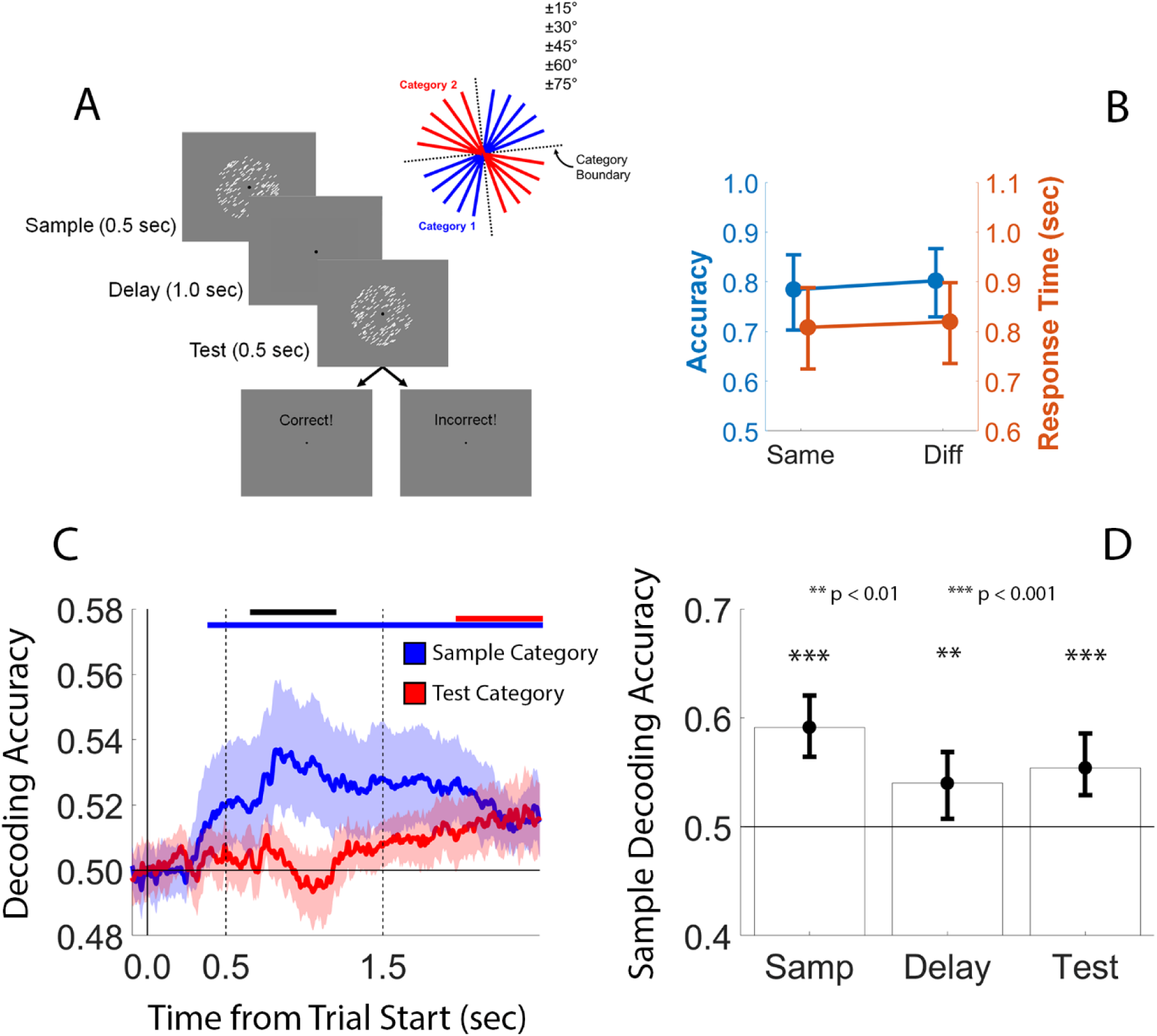
Results of Experiment 2. (A) Volunteers performed a delayed match-to-category (DMC) task where they judged whether two successively presented stimuli (the sample and test) were drawn from the same category. (B) Behavioral performance during same and different trials. (C) A linear discriminant function trained to classify the sample stimulus’ category supported robust above-chance decoding accuracy ∼0.4 sec after sample onset and throughout the subsequent delay and test periods. A separate decoder trained to classify the test stimulus’ category also supported robust above-chance decoding accuracy ∼0.4 sec after test onset. The solid vertical line at time 0.0 depicts the onset of the sample stimulus, while the dashed vertical lines at time 0.5 and 1.5 depict the onset of the delay and test periods. (D) Support vector machine classifiers trained on fixations measured during the sample, delay, and test periods supported above-chance decoding of the sample category. Shaded regions and error bars in panels B-D depict the 95% confidence interval of the mean; the horizontal solid line at 0.5 in panels C and D depicts chance decoding accuracy. The blue bar and red bars at the top of panel C depicts epochs where sample and test category decoding accuracy was significantly above chance (nonparametric signed randomization test; see Methods). The black bar depicts epochs where sample decoding accuracy was significantly higher than test

#### Experiment 3

##### Data Exclusion

Data from 3 of the 16 volunteers who completed Experiment 3 were discarded. Two volunteers withdrew from the study, and data from one volunteer were discarded due to hardware faults that occurred mid-testing. Thus, the data reported here reflect the remaining 13 volunteers.

##### Experimental Tasks

Volunteers performed two tasks that used identical stimuli (Figure 5A-B). Volunteers viewed displays containing flickering apertures of iso-oriented black and white bars. The proportion of white bars varied across trials from 15% to 85% in 10% steps. During the color task, participants reported whether the display contained a greater proportion of white vs. black bars. Color category was defined according to this majority: displays with >50% white bars were assigned to one color category, and displays with >50% black bars were assigned to the other.

**Figure 5.**
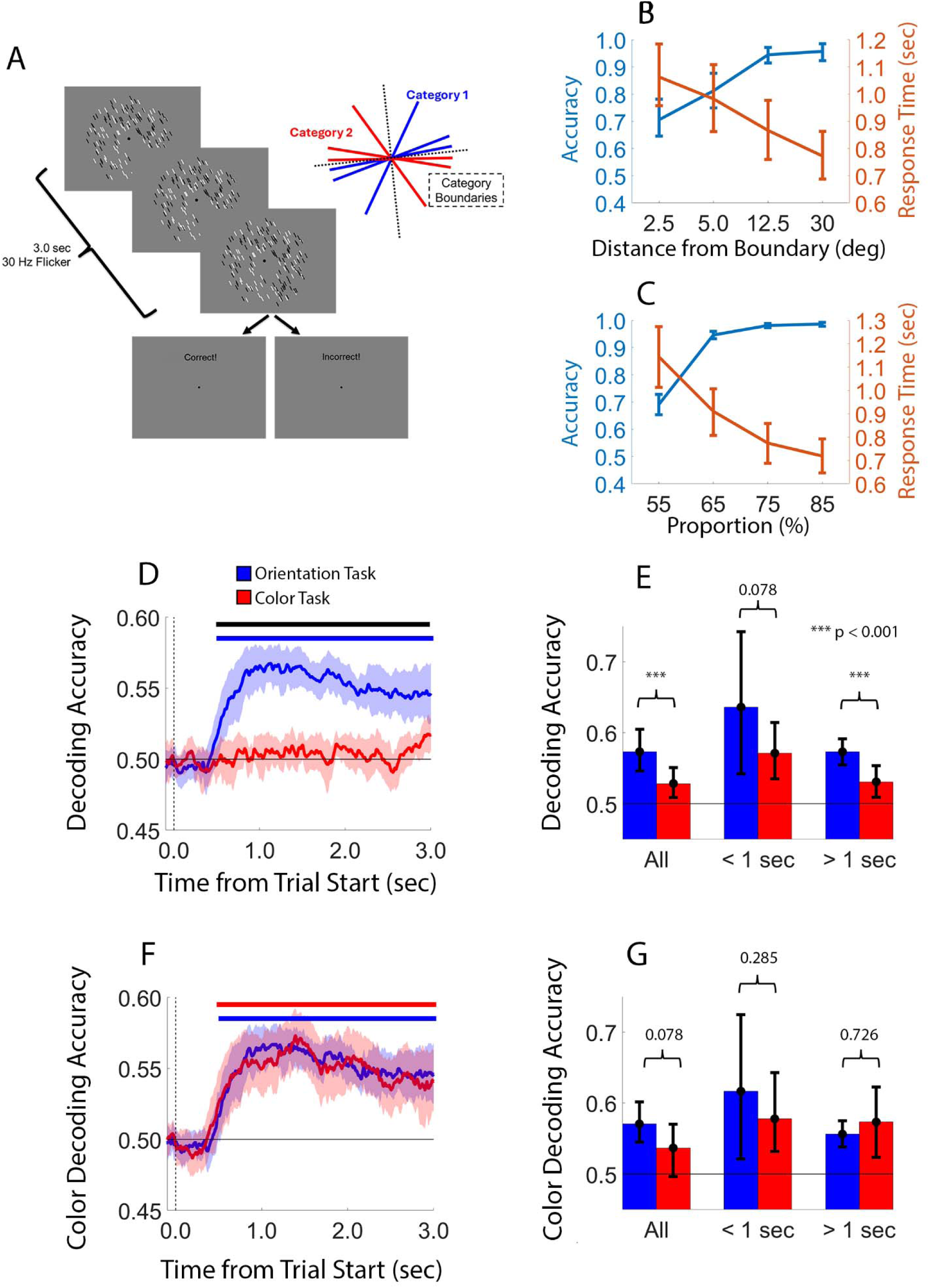
Results of Experiment 3. (A) Volunteers viewed circular apertures containing flickering black and white bars while performing two tasks with identical stimuli. In the first half of the experiment, they classified stimulus orientation relative to a memorized boundary (orientation task; inset). In the second half, they judged whether each display contained a greater proportion of white or black bars (color task). (B) Accuracy and response times during the orientation task. (C) Accuracy and response times during the color task. (D) Orientation-category decoding accuracy during the orientation and color tasks. The blue bar indicates epochs where decoding exceeded chance, and the black bar indicates epochs where decoding was greater in the orientation than the color task. Support vector machine classifiers trained on fixations from the full stimulus interval (“all”), the first second (“< 1 s”), and the final two seconds (“> 1 s”) yielded above-chance orientation-category decoding in both tasks, with generally higher accuracy in the orientation task. (E) Orientation-category decoding from stable fixations restricted to the stimulus aperture. (F) Color-category decoding (majority white vs. black bars) from fixation patterns during the orientation and color tasks; decoding was robust and similar in magnitude across tasks. (G) Color-category decoding from stable fixations, showing reliable and comparable above-chance performance in both tasks. Shaded regions and error bars depict 95% confidence intervals of the mean.

During the first half of the experiment, volunteers performed a categorization task that was nearly identical to Experiment 1. We defined a set of oriented stimuli ±2.5°, ±5.0°, ±12.5°, and ±30° relative to the volunteer’s category boundary, with negative values assigned to Category 1 and positive values assigned to Category 2. Volunteers reported the category membership of presented stimuli by pressing the “z” and “?” keys (for Categories 1 and 2, respectively) at any point during each three-second trial. Failures to respond within the 3 second stimulus window were treated as incorrect. Volunteers completed 7 (N = 8) or 8 (N = 5) blocks of 64 trials.

During the second half of the experiment, volunteers viewed physically identical stimuli but instead reported whether they contained a greater proportion of white vs. black bars regardless of orientation category membership, pressing “z” to indicate a greater proportion of white bars and “?” to indicate a greater proportion of black bars. Failures to respond within the 3 second stimulus window were treated as incorrect. Volunteers completed 4 (N = 3), 6 (N = 1), 7 (N = 6), or 8 (N = 3) blocks of 64 trials.

## Results

### Training Tasks

All volunteers completed a training regimen before starting the main experimental tasks. For each volunteer and each experiment, we randomly selected a category boundary on the interval (0°,179°] and defined stimuli with orientations ±15°, ±30°, ±45°, and ±60° (Experiments 1 and 3; Figure S1) or ±15°, ±30°, ±45°, ±60° and ±75° relative to the boundary (Experiment 2). Stimuli with negative values (i.e., counterclockwise from the boundary) were assigned to Category 1, while stimuli with positive values (i.e., clockwise from the boundary) were assigned to Category 2. Volunteers learned to categorize stimuli through trial-and-error, with feedback given at the end of each trial. Volunteers learned to categorize stimuli quickly, frequently reaching asymptotic performance in 1-2 blocks of 32 trials (Figure S1). Each volunteer’s unique category boundary was used in subsequent experimental tasks.

### Experiment 1

Volunteers in Experiment 1 classified oriented stimuli into discrete groups based on a learned boundary, providing a behavioral benchmark for subsequent gaze-based decoding analyses (Figure 1A). To make the categorization task more difficult, and to ensure that volunteers classified stimuli by rule vs. rote memory, we defined a stimulus set containing some orientation values volunteers had not seen during training (±0, ±2°, ±5°, ±15°, ±45°; Figure 1B). As expected, volunteers’ accuracy increased, and their response times decreased, with angular distance from the category boundary. We found no differences in behavioral performance between stimuli tilted counterclockwise vs. clockwise relative to the category boundary, therefore, we labeled conditions according to their absolute angular distance from the boundary (Figure 1C).

#### Eye Movements Reflect Abstract Categories

We first asked whether trial-wise gaze position could be used to decode stimulus category during the speeded classification task. Specifically, we tested whether volunteers’ eye movements varied with stimulus category by training a linear classifier to predict category membership from time-resolved patterns of Cartesian gaze position. This analysis omitted data from ambiguous trials (i.e., ±0°); we return to these trials below. Eye position data from N-1 experimental blocks were used to train the classifier, and the trained classifier was used to predict category membership from data in the held-out block. This procedure was repeated until every experimental block had served as the test data set. This analysis was performed on a timepoint-by-timepoint basis, yielding a single decoding time course per volunteer. Since there are only two categories, observed decoding accuracy was evaluated relative to a chance value of 50%. This analysis revealed modest but robust above-chance decoding of category membership, mostly within an interval spanning ∼0.7 to ∼1.4 seconds after stimulus onset (Figure 2A). Note that this interval corresponds to the period during which most behavioral responses occurred (Figure 1C). To better visualize relationships between decoding accuracy and behavioral responses, we re-computed decoding accuracy over an interval from -1.0 to +0.5 sec around volunteers’ responses. Decoding accuracy was stable during this period, but variability in decoding accuracy decreased near the time of a behavioral response (Figure 2B).

Because time-resolved category decoding accuracy could be jointly influenced by fixations and saccades, we re-computed category decoding accuracy using stable fixations (i.e., saccade endpoints). Specifically, we trained linear support vector machines to classify stimulus category based on the cartesian coordinates of fixations measured at any point during each 3 second trial (“all”), those measured before a response time threshold, defined as the 80^th^ percentile of each volunteer’s response time distribution across all trials and conditions (“< RT”), and those measured after the response time threshold (“> RT”). Fixations were identified using a velocity-based threshold procedure (see Methods), and we discarded fixations that fell outside the 9 DVA stimulus aperture. This analysis revealed robust decoding of stimulus category (Figure 2C; p < 1e-04 for the “all”, “< RT”, and “> RT” epochs; bootstrap tests). Identical results were obtained when we restricted this analysis to fixations inside the inner radius of the stimulus aperture (2 DVA; Figure S2A).

We also asked whether category decoding performance varied with the accuracy of participants’ category judgments. Although our data set contained too few incorrect trials to enable direct comparisons, we did have sufficient data to undertake a comparison of category decoding accuracy using classifiers trained using data from all trials vs. only correct trials. This analysis revealed modestly but significantly greater category decoding accuracy during correct trials (Figure 2D), indicating that category-linked differences in eye movements are more pronounced when the categorical decision is correct, consistent with the idea that gaze patterns reflect the internal decision state rather than only the presented stimulus. We observed similar, though less robust results when we trained classifiers on fixations within the inner radius of the stimulus aperture (2 DVA, respectively; Figure S2B).

We also considered the possibility that category decoding was driven by stimulus-evoked eye movements rather than abstract categorical representations. For example, eye movements might be systematically anchored to the physical orientation of the stimulus relative to the boundary (counterclockwise vs clockwise), yielding a continuous gradient of oculomotor biases that a classifier could exploit. If this were the case, decoding accuracy should increase monotonically with the angular distance between the stimulus and the boundary, because larger distances correspond to greater physical distinctiveness between categories. To test this prediction, we trained a support vector machine to distinguish stimulus category using all non-ambiguous trials (±2°, ±5°, ±15°, ±45°) and then applied the trained classifier to held-out data sorted by angular distance from the boundary. Decoding performance was statistically equivalent across all four distances (Figure 2E; F(3, 81) = 1.490, p = .223, η² = 0.05), yielding a flat profile. This pattern is inconsistent with a purely stimulus-driven gradient account and instead supports the interpretation that the classifier detected a discrete, category-selective signal in gaze behavior.

#### Category Decoding Depends on Multiple Eye Movement Parameters

Next, we sought to determine the eye movement parameter(s) responsible for above-chance category decoding accuracy. Visual inspection of volunteers’ fixations revealed no obvious category-specific patterns (Figure 3A). We first tested whether decoding accuracy was driven by differences in the number of saccades recorded during Category 1 or Category 2 trials (irrespective of saccade amplitude or direction). Although saccade frequencies varied substantially across volunteers (e.g., from a few hundred to a few thousand), we found no evidence for category level-differences in saccade frequency (Figure 3B). Similarly, total saccade count was uncorrelated with category decoding accuracy (Figure S3A). Next, we considered whether decoding was driven by category-level differences in saccade size (irrespective of frequency and direction). Most saccades were small (e.g., within 2-3 DVA of fixation) and their amplitudes were remarkably similar across categories (Figure 3C). Similarly, individual differences in average saccade size were uncorrelated with category decoding accuracy (Figure S3B). Finally, we tested whether decoding accuracy was driven by category-level differences in saccade direction. For this analysis, we aligned volunteers’ fixation records to a common orientation (0°; note that this step was necessary because each volunteer classified stimuli according to a bespoke category boundary) and constructed circular histograms plotting gaze frequency across each of 360 bins (i.e., 1° bin widths). Saccade directions were remarkably similar and statistically uniform during both Category 1 and Category 2 trials (Figure 3D). Two-sample Kuiper tests (see Methods) used to evaluate category-level differences in fixation coordinates revealed a significant outcome in only 5 of 28 volunteers. Thus, although linear classifiers could successfully decode category membership from records of volunteers’ eye movements, we could not identify a single eye movement parameter responsible for these differences.

To quantify links between specific eye movement parameters and category membership, we fit a mixed-effects logistic regression with stimulus category as the dependent variable and four eye movement parameters as fixed-effect predictors: saccade rate, mean saccade amplitude, mean horizontal saccade direction (cosine-transformed), and mean vertical saccade direction (sine-transformed), with random intercepts for participants (see Methods). Neither rate (β = −0.0285, *p* > .999) nor horizontal direction (β = −0.0789, *p* = .92) predicted stimulus category. In contrast, vertical direction (β = 0.0821, *p* < .001) and amplitude (β = 0.0005, *p* = .01) were reliable, albeit weak, predictors (Table 1). Although these individual effects were small, their presence suggests that category information is distributed across multiple oculomotor parameters rather than concentrated in any single gaze feature, a pattern consistent with the phasic and idiosyncratic category-selective eye movements reported in non-human primates performing analogous tasks (Rosen & Freedman, 2025a).

#### Motor Confounds Do Not Account for Category Decoding

The design of Experiment 1 confounded category membership with motor responses: volunteers always pressed “Z” for Category 1 and “?” for Category 2. Thus, category decoding could reflect gaze shifts toward the spatially separated response keys on the keyboard rather than abstract category representations. We tested this possibility using ambiguous trials—those where the presented orientation precisely matched the category boundary (±0°)—which contain no category information and yet still require a motor response. We trained a linear classifier on gaze data from unambiguous trials (±2°, ±5°, ±15°, ±45°), then tested whether the trained classifier could predict volunteers’ responses on ambiguous trials. If category decoding reflects eye movements toward the response key selected on a given trial, the classifier should successfully predict these responses. However, this analysis failed: decoding accuracy on ambiguous trials did not differ from chance (Figure S4). This null result is difficult to reconcile with a response-key gaze account, because on ambiguous trials volunteers still selected and executed a key press, but the classifier trained on unambiguous trials could not recover their choice from gaze. Nevertheless, because category and response remained confounded at the design level in Experiment 1, we turned to Experiment 2 to more stringently dissociate category encoding from response preparation.

### Experiment 2

Experiment 2 asked whether category-selective gaze patterns emerge when category and response are temporally decoupled, using a delayed match-to-category task in which the correct response is undefined until the test stimulus appears (Freedman & Assad, 2006). On each trial, volunteers viewed a sample stimulus, a blank delay, and a test stimulus, and reported whether the sample and test belonged to the same category (Figure 4A–B). Critically, the correct response (same vs. different) depended on the pairing of sample and test categories and was therefore undefined until the test appeared. Any category-selective gaze patterns during the sample (0–500 ms) or delay (500–1500 ms) periods thus cannot reflect response selection or motor preparation.

Behaviorally, volunteers performed the DMC task with high accuracy and reaction times that were well separated from the sample and delay periods (Figure 4B), confirming that participants understood and used the category rule. We first tested whether gaze position during the sample and delay periods carried information about the category of the sample stimulus. A linear discriminant classifier trained to distinguish the sample category supported robust above-chance decoding beginning approximately 0.4 s after sample onset and persisting throughout the delay and test periods (Figure 4C). A separate decoder trained on the test category likewise yielded above-chance decoding beginning ∼0.4 s after test onset (Figure 4C). Sample-based decoding was significantly higher than test-based decoding during the delay period, consistent with the idea that gaze patterns during this interval primarily reflect the category of the remembered sample rather than anticipatory responses to the yet-to-be-seen test. Because no response could be prepared during the sample and delay periods, these results demonstrate that time-resolved category-selective gaze patterns cannot be explained by simple eye movements toward the eventual response.

To summarize category information in distinct processing stages, we next computed epoch-based decoding accuracy from fixations during the sample, delay, and test intervals. Support vector machine classifiers trained on fixations recorded during each epoch supported reliable above-chance decoding of the sample category across all three periods (Figure 4D). Together with the time-resolved decoding, these fixation-based analyses indicate that subtle differences in gaze position corresponding to stimulus category persist throughout the trial, including periods when no motor plan can be specified. The presence of category information in gaze during the delay period again suggests that eye movements track an internal categorical representation maintained across time, not just immediate sensory input or imminent motor output.

Finally, we again used mixed-effects logistic regression to identify which eye-movement parameters contributed to category decoding in the DMC task. We fit a mixed-effects logistic regression model, analogous to Experiment 1, with sample category (Category 1 vs. Category 2) as the dependent variable and saccade rate, mean saccade amplitude, and mean horizontal and vertical saccade directions as predictors. In contrast to Experiment 1, where vertical direction and amplitude provided small but significant prediction, saccade rate (number of saccades per trial) emerged as the only reliable predictor of category membership in Experiment 2, whereas amplitude and horizontal and vertical directions were not significant (all p ≥ 0.695; Table 2). This pattern reinforces the conclusion that category information cannot be consistently localized to a single gaze parameter and instead is expressed via weak, task-dependent combinations of oculomotor features.

Together, the Experiment 2 results show that human gaze behavior tracks the category of a remembered sample even when the appropriate response is not yet defined, ruling out simple motor-preparation accounts of gaze-based decoding. We next asked whether these category-selective gaze patterns depend on the current task context or instead reflect fixed stimulus-driven biases, by comparing decoding across orientation- and color-judgment tasks with identical visual input (Experiment 3).

### Experiment 3

The designs of Experiments 1 and 2 confounded orientation values with category labels: different orientations were always assigned to different categories. Thus, what we have described as category-selective gaze patterns could, in principle, reflect orientation-specific eye movements rather than abstract categorization. Experiment 3 tested this possibility using two tasks that employed identical oriented stimuli but required different judgments. During the first half of each block, volunteers performed an orientation categorization task that was nearly identical to Experiment 1 (orientation task). During the second half, they viewed the same displays but instead judged whether the aperture contained a greater proportion of white versus black bars (color task), which required attentive processing but no orientation category judgment (Figure 5A).

Volunteers performed both the orientation and color tasks with high accuracy, confirming that they successfully learned and applied the respective decision rules (Figure 5B–C). Response times were comparable across tasks, indicating similar overall difficulty and engagement. Due to a coding error, response time data during the orientation and color tasks for one volunteer were irretrievably lost; the RT analyses therefore include 12 volunteers, whereas other analyses include all 13.

We first asked whether gaze position distinguished orientation categories during the orientation task, replicating the main finding from Experiment 1 with physically identical stimuli used across tasks. Using fixation-based decoding restricted to the stimulus aperture, we observed robust above-chance decoding of orientation category during the orientation task (Figure 5D). Thus, when observers were explicitly categorizing orientation, gaze patterns again carried information about the rule-defined orientation categories.

To summarize orientation information across different processing stages, we computed decoding accuracy from fixations measured (1) at any point during the 3-s stimulus window, (2) during the first second of each trial (when most responses occurred), and (3) during the final 2 seconds (after most responses were complete). Orientation category could be decoded from fixations during the orientation task in all three epochs. Across epochs, decoding accuracy was generally higher in the orientation than in the color task (Figure 5E). These results indicate that the two tasks evoke different patterns of eye movements with respect to orientation category, even though they use a common set of oriented stimuli, and that robust orientation-based category signals in gaze depend on task demands.

We next tested whether gaze patterns carried information about color category, defined by the majority of white versus black bars in each display. During the color task, color category could be reliably decoded from fixation locations (Figure 5F-G), indicating that gaze behavior tracks the currently relevant color decision. Strikingly, color category was also decodable during the orientation task, despite color being irrelevant to the observer’s explicit judgment. This suggests that oculomotor behavior can simultaneously reflect both the task-relevant orientation category and the task-irrelevant color category when both features covary with the visual input.

Taken together, these within-task analyses reveal a partial dissociation: during orientation categorization, gaze behavior carries information about both orientation and color categories, whereas during color categorization, gaze primarily reflects color category and only weak or absent orientation category information. This asymmetry suggests that orientation-based category signals in gaze may depend more strongly on task set and decision demands than do color-based signals, which appear whenever color is behaviorally relevant and even when it is not.

Finally, we implemented cross-task decoding analyses to test the extent to which the orientation and color tasks evoked shared category-linked gaze patterns. In the first phase of this analysis, we trained a linear discriminant classifier to predict orientation category using continuous gaze position from the orientation task, then asked whether the trained classifier could decode orientation category from gaze during the color task. This analysis failed to reveal above-chance decoding at any point during the 3-s stimulus interval (Figure 6). In a second phase, we trained a support vector machine to decode orientation category from fixations measured at any point during the stimulus interval in the orientation task and applied it to fixations from the color task; again, cross-task decoding was at chance (M = 0.482, p = 0.856, bootstrap test).

**Figure 6.**
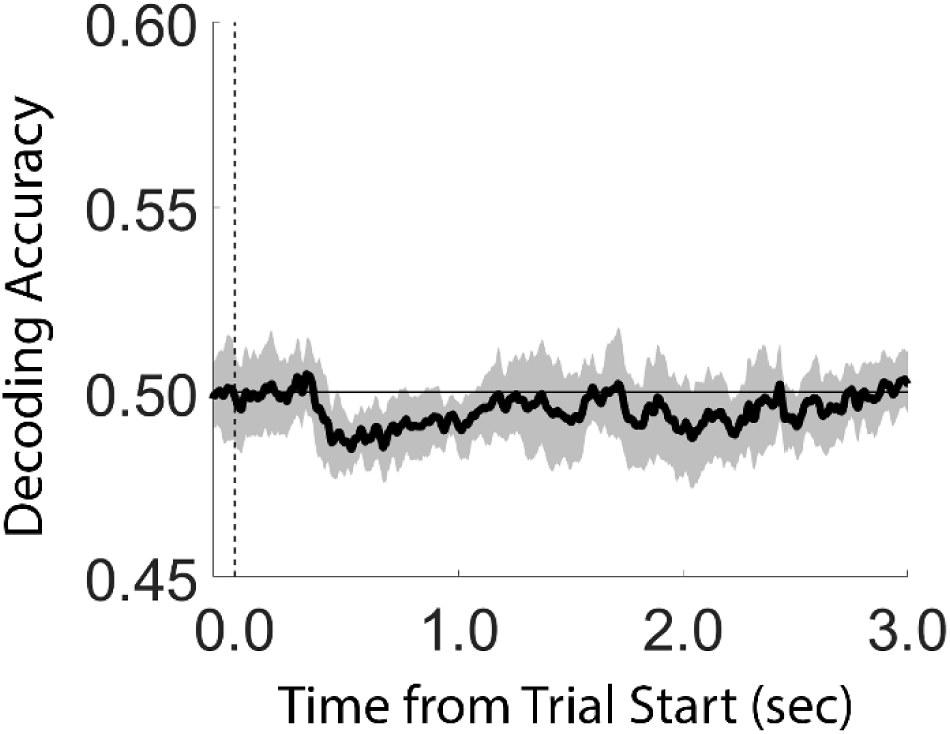
Cross-Task Orientation Category Decoding Performance. A linear classifier was trained to predict orientation category (Category 1 vs. Category 2) from fixation patterns during the orientation task in Experiment 3, then applied to data from the color task, which used identical oriented stimuli but required a color majority judgment. Cross-task decoding remained at chance throughout the trial, indicating that orientation-and color-task trials evoked distinct patterns of eye movements despite a common stimulus set and identical response keys.

Thus, although orientation category could be decoded from gaze in the orientation task, the patterns of eye movements that supported orientation category decoding did not generalize to the color task. Because the two tasks used identical oriented stimuli, this lack of crossl1ltask generalization indicates that orientation category decoding cannot be attributed to fixed, orientationl1lspecific eye movements and instead reflects taskl1ldependent categorical processing.

## Discussion

The present findings demonstrate that human gaze behavior carries reliable information about abstract, rule-defined stimulus categories. Across three experiments, category membership could be decoded from trial-wise gaze position during speeded classification (Experiment 1), during a delayed match-to-category task in which the correct response was undefined until the test stimulus appeared (Experiment 2), and across tasks that used identical stimuli but imposed different decision rules (Experiment 3). Category-selective gaze patterns were stronger on correct trials, present during delay periods when no response could yet be prepared, and shaped by task demands and feature relevance. Together, these results indicate that incidental eye movements reflect the observer’s internal categorical decision state rather than only the physical properties of the stimulus or overt motor responses.

These behavioral results converge with neurophysiological evidence linking abstract cognition to oculomotor circuits. In macaques, superior colliculus neurons encode learned visual categories with short latency and strength that can exceed posterior parietal cortex, and SC inactivation selectively impairs categorization (Peysakhovich et al., 2024). Rosen and Freedman (2025a) showed that small, uninstructed eye movements during delayed match-to-category tasks reflect the sample’s category and arise from transient alignment between cognitive and saccadic coding subspaces in SC—shifts that were absent under non-categorical task demands, implicating a multiplexing mechanism that embeds decision variables within motor dynamics. Our human data provide a behavioral analogue: category-selective eye movements appear during rule-based categorization, are modest in magnitude, and are distributed across multiple gaze parameters rather than concentrated in any single oculomotor feature.

Several aspects of these results are consistent with the proposed orthogonal-subspace mechanism for cognitive–motor multiplexing. First, the decoded category signal is small but robust, consistent with partial rather than complete alignment between cognitive and saccadic codes: category information projects weakly onto individual gaze dimensions yet remains recoverable with multivariate decoding. Second, mixed-effects regression revealed that different oculomotor parameters predicted category across experiments—vertical direction and amplitude in Experiment 1, saccade rate alone in Experiment 2—indicating that category information is distributed across gaze features rather than tied to a single stereotyped movement pattern. This parallels observations in monkeys that category-linked eye movement parameters vary idiosyncratically across animals and sessions (Rosen & Freedman, 2025a), precisely the pattern expected if the category signal reflects a low-dimensional projection from a high-dimensional cognitive subspace onto a differently oriented motor subspace. Third, category-selective gaze patterns in both species are task-dependent, appearing during rule-based categorization and absent or attenuated under different task demands (Experiment 3). This diffuse, task-dependent encoding profile explains why univariate analyses of individual gaze parameters yield weak or null effects while multivariate decoding succeeds, and suggests a conserved mechanism by which abstract task variables modulate primate oculomotor dynamics without disrupting primary orienting functions.

Our findings also extend prior non-human primate work in several important ways. First, whereas most SC categorization studies have used extrafoveal motion stimuli that engage spatially tuned oculomotor neurons, our stimuli were foveally presented, and category decoding did not vary with angular distance from the category boundary (Experiment 1). This flat distance-to-boundary profile is incompatible with a stimulus-driven gradient and indicates that spatial coding is not required to observe category-selective gaze patterns. Second, category-modulated gaze emerged rapidly in humans: participants reached asymptotic performance within a few dozen trials, in contrast to the extensive training regimes used in monkeys (Freedman et al., 2001; Peysakhovich et al., 2024), raising the possibility that explicit rule learning and frontoparietal control systems may accelerate the recruitment of oculomotor circuits for abstract decisions. Third, gaze patterns encoded category information about color—a non-spatial feature that should not, by itself, impose a directional eye movement bias—with robust color-based decoding present in both orientation and color tasks (Experiment 3). This feature-general expression of categorical information addresses open questions about whether oculomotor involvement in categorization is specific to spatial features and suggests that higher-order category signals can modulate eye movements even when categories are defined along non-spatial dimensions.

Our approach also complements recent evidence that microsaccade rate distributions can discriminate natural object categories during passive viewing (Nouri et al., 2024). That study decoded ecological categories (e.g., animate vs. inanimate) from temporal microsaccade profiles, achieving high classification accuracy in both monkeys and humans. The present findings extend this work in three respects: the categories decoded here were abstract and rule-defined rather than ecologically grounded, decoding relied on spatial gaze position rather than temporal rate profiles, and a task-dependence manipulation (Experiment 3) established that the decoded signal reflects the observer’s current decision state—a dimension that passive viewing paradigms cannot address. More broadly, these results invert a concern raised by prior work showing that task-specific eye movements can confound neural decoding of working memory representations (Mostert et al., 2018; Quax et al., 2019): whereas those studies identified eye movements as a nuisance variable that contaminates neural signals, our findings—together with those of Nouri et al. (2024) and Rosen and Freedman (2025a)—suggest that these same eye movements carry genuine cognitive information and can serve as a direct, non-invasive readout of internal categorical states.

The rule-based paradigm used here captures only one form of categorization. Humans and other animals also learn categories defined by multidimensional feature combinations (Goldstone, 1994; Reinert et al., 2021; Henderson et al., 2025) that rely more heavily on procedural learning in the striatum and basal ganglia (Ashby & Maddox, 2005) than on the frontoparietal networks recruited for rule-based categorization. Whether these distinct learning systems produce comparable oculomotor signatures is unknown. Given established basal ganglia–SC connectivity (Melleu & Sabino Canteras, 2024), it is plausible that categorical information can modulate oculomotor control regardless of learning format; alternatively, gaze modulation may be specific to rule-based tasks that engage frontoparietal cognitive control, consistent with the task-dependence of SC category encoding observed by Rosen and Freedman (2025a). Future work directly comparing rule-based and information-integration tasks could clarify how different learning systems shape the coupling between decision representations and oculomotor dynamics.

Our findings also dovetail with recent proposals that working memory and decision variables are implemented via flexible “sensory reformatting” rather than static sensory traces (Kiyonaga & Serences, 2025). Gaze patterns in the present study did not simply mirror low-level stimulus features but expressed category information in a way that depended on task set, temporal epoch, and the relevance of different feature dimensions. In this view, the category-selective structure we observe in gaze reflects one manifestation of a broader process by which distributed sensorimotor circuits are recruited and reformatted to carry abstract, task-dependent cognitive codes.

At a broader level, the present findings speak to an emerging theme in systems neuroscience: that cognitive signals are far more widely distributed across brain circuits—including subcortical and motor structures—than traditionally appreciated (Rosen & Freedman, 2025b; Peysakhovich et al., 2024). The oculomotor system, long studied as an effector for spatial orienting, now appears to be a reliable window onto abstract cognitive states, from the spatial coordinates of remembered stimuli (van Ede et al., 2019) to their categorical identity (present study). That these signals appear in both human and non-human primate gaze—despite vast differences in training, stimuli, and response modality—suggests a conserved computational architecture in which abstract task variables are multiplexed within oculomotor networks. Determining which task demands recruit oculomotor circuits, which stimulus features modulate the gaze signal, and how learning reshapes cognitive–motor coupling represents a rich direction for future work at the intersection of systems, cognitive, and computational neuroscience.

## Author Contributions (CRedIT)

Conceptualization: EFE, AC

Methodology: EFE, AC

Software: EFE

Validation: AC, EFE

Formal Analysis: EFE, AC

Investigation: AC

Resources: EFE

Data Curation: EFE, AC

Writing – Original EFE

Writing – Editing AC, EFE

Visualization: AC, EFE

Supervision: EFE

Project Administration EFE

Funding Acquisition EFE

## Acknowledgements

We thank Ms. Jasdeep Buttar for assistance with data collection.

## References

Ashby, F. G., & Maddox, W. T. (2005). Human Category Learning. Annual Review of Psychology, 56(1), 149–178. 10.1146/annurev.psych.56.091103.070217

Chota, S., Arora, K., Kenemans, J. L., Gayet, S., & Van Der Stigchel, S. (2025). Microsaccade biases can reflect task-specific spatial memorization strategies. Frontiers in Neuroscience, 19, 1526213. 10.3389/fnins.2025.1526213

de Vries, E., van Ede, F. (2024) Microsaccades track location-based object rehearsal in visual working memory. eNeuro, 11, 10.1523/ENEURO.0276-23.2023

Ester, E. F., Sprague, T. C., & Serences, J. T. (2020). Categorical Biases in Human Occipitoparietal Cortex. The Journal of Neuroscience, 40(4), 917–931. 10.1523/JNEUROSCI.2700-19.2019

Ester, E., & Weese, R. (2023). Temporally Dissociable Mechanisms of Spatial, Feature, and Motor Selection during Working Memory–guided Behavior. Journal of Cognitive Neuroscience, 35(12), 2014–2027. 10.1162/jocn_a_02061

Freedman, D. J., Riesenhuber, M., Poggio, T., & Miller, E. K. (2001). Categorical Representation of Visual Stimuli in the Primate Prefrontal Cortex. Science, 291(5502), 312–316. 10.1126/science.291.5502.312

Freedman, D. J., & Assad, J. A. (2006) Experience-dependent representation of visual categories in the parietal cortex. Nature, 443, 85–88.

Goldstone, R. L. (1994). Influences of categorization on perceptual discrimination. Journal of Experimental Psychology: General, 123(2), 178–200. 10.1037/0096-3445.123.2.178

Henderson, M. M., Serences, J. T., & Rungratsameetaweemana, N. (2023). Dynamic categorization rules alter representations in human visual cortex. 10.1101/2023.09.11.557257

Kiyonaga, A., & Serences, J. T. (2025) Sensory reformatting for a working visual memory. Trends Cogn Sci. 29(12) 1120–1135.

Linde-Domingo, J., & Spitzer, B. (2023). Geometry of visuospatial working memory information in miniature gaze patterns. Nature Human Behaviour, 8(2), 336–348. 10.1038/s41562-023-01737-z

Liu, B., Alexopoulou, Z.-S., & Van Ede, F. (2024). Jointly looking to the past and the future in visual working memory. eLife, 12, RP90874. 10.7554/eLife.90874

Maris, E., & Oostenveld, R. (2007). Nonparametric statistical testing of EEG- and MEG-data. Journal of Neuroscience Methods, 164(1), 177–190. 10.1016/j.jneumeth.2007.03.024

Melleu, F. F., & Sabino Canteras, N. (2024) Pathways from the superior colliculus to the basal ganglia. Current Neuropharmacology, 22, 1431–1453. 10.2174/1570159X21666230911102118

Mostert, P., Albers, A. M., Brinkman, L., Todorova, L., Kok, P., & de Lange, F. P. (2018). Eye movement-related confounds in neural decoding of visual working memory representations. eNeuro, 5(4), ENEURO.0401 17.2018. 10.1523/ENEURO.0401-17.2018

Nouri, S., Soltani Tehrani, A., Faridani, N., Toosi, R., Noroozi, J., & Dehaqani, M.-R. A. (2024). Microsaccade selectivity as discriminative feature for object decoding. iScience, 28(1), 111584. 10.1016/j.isci.2024.111584

Peysakhovich, B., Zhu, O., Tetrick, S. M., Shirhatti, V., Silva, A. A., Li, S., Ibos, G., Rosen, M. C., Johnston, W. J., & Freedman, D. J. (2024). Primate superior colliculus is causally engaged in abstract higher-order cognition. Nature Neuroscience. 10.1038/s41593-024-01744-x

Quax, S. C., Dijkstra, N., van Staveren, M. J., Bosch, S. E., & van Gerven, M. A. J. (2019). Eye movements explain decodability during perception and cued attention in MEG. NeuroImage, 195, 444–453. 10.1016/j.neuroimage.2019.03.069

Reinert, S., Hübener, M., Bonhoeffer, T., & Goltstein, P. M. (2021) Mouse prefrontal cortex represents learned rules for categorization. Nature, 593, 411–417. 10.1038/s41586-021-03452-z

Rosen, M. C., & Freedman, D. J. (2025a) Multiplexing of cognitive encoding by oculomotor networks leads to incidental gaze shifts. Proceedings of the National Academy of Sciences USA 122 10.1073/pnas.2422331122

Rosen, M. C., & Freedman, D. J. (2025b) How distributed is the brain-wide network that is recruited for cognition ? Nature Reviews Neuroscience, 27, 138–150 10.1038/s41583-025-00992-5

Swaminathan, S. K., & Freedman, D. J. (2012) Preferential encoding of visual categories in parietal cortex compared with prefrontal cortex. Nature Neuroscience, 15, 315–320. 10.1038/nn.3016

van Ede, F., Chekroud, S. R., & Nobre, A. C. (2019). Human gaze tracks attentional focusing in memorized visual space. Nature Human Behaviour, 3(5), 462–470. 10.1038/s41562-019-0549-y

Zhou, Y., & Freedman, D. J. (2019). Posterior parietal cortex plays a causal role in perceptual and categorical decisions. Science, 365(6449), 180–185. 10.1126/science.aaw8347

Zhou, Y., Zhu, O., & Freedman, D. J. (2023). Posterior Parietal Cortex Plays a Causal Role in Abstract Memory-Based Visual Categorical Decisions. The Journal of Neuroscience, 43(23), 4315–4328. 10.1523/JNEUROSCI.2241-22.2023

