## Supplemental Figures for "Human gaze behaviors track abstract stimulus categories"

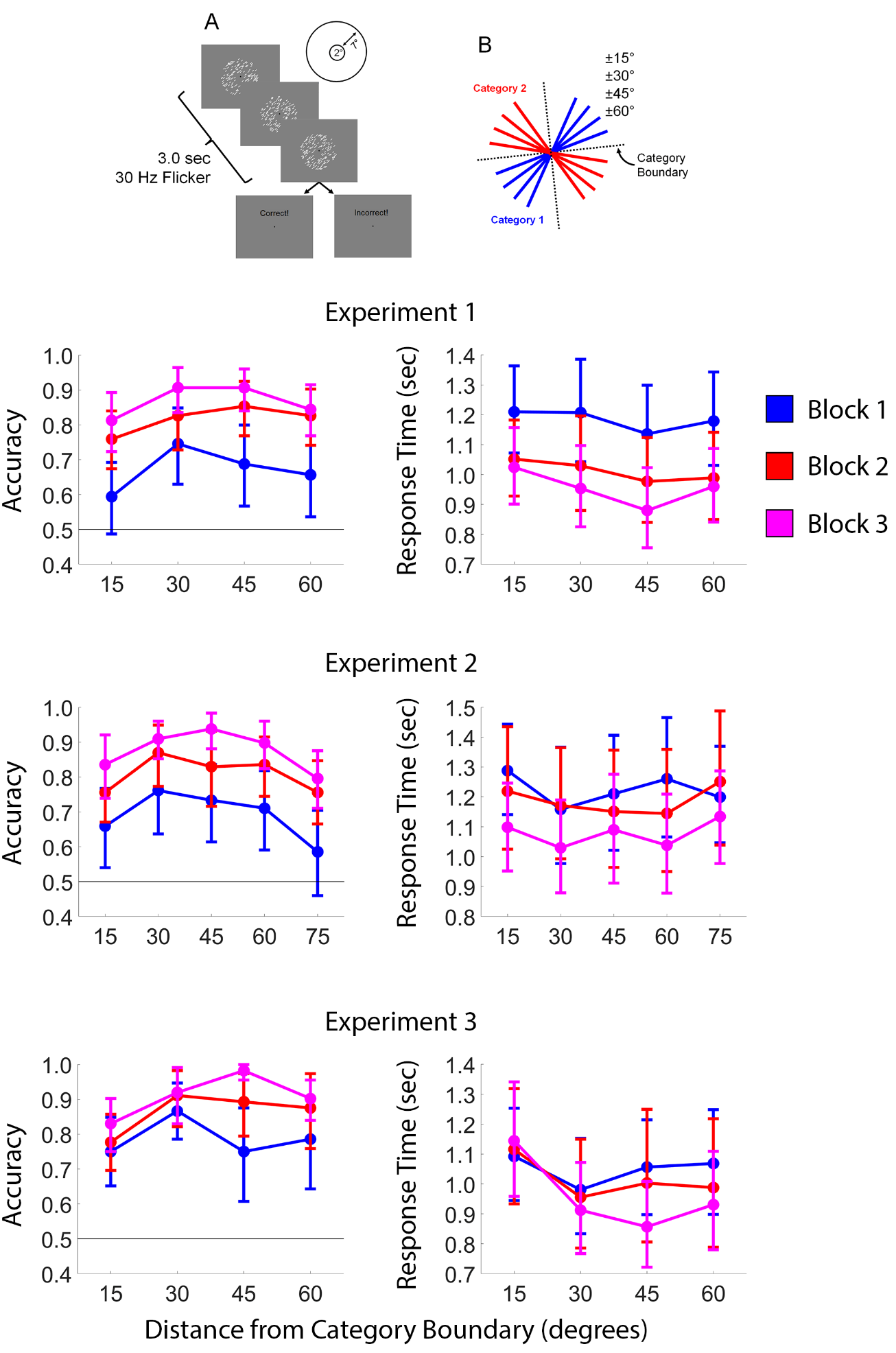

**Figure S1. Training Data.** (A,B) Participants in each experiment completed 3 blocks of 32 trials in a training task. Participants learned to classify sets of continuously oriented stimuli into discrete groups according to a category boundary through trial-and-error. Participants accuracies increased, and response times decreased, across blocks 1-3. Most participants reached asymptotic task performance in 2-3 blocks. Error bars depict the 95% confidence interval of the mean.

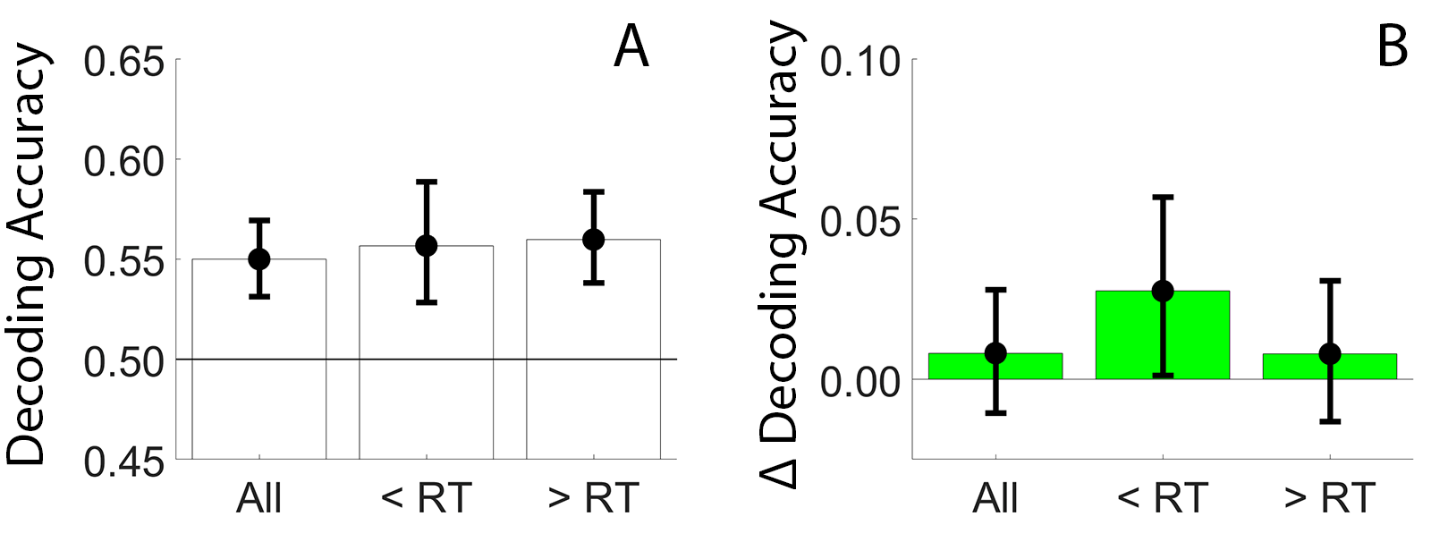

**Figure S2. Decoding Accuracy During Experiment 1 Calculated Using Microsaccades.** (A) Decoding accuracy calculated using small eye movements (< 2 DVA) was similar to performance using all recorded eye movements (see Figure 2C). (B) Change in category decoding performance computed during correct vs. all trials. Category decoding performance during the pre-response time threshold (“< RT”) was significantly greater during correct vs. all trials (p = 0.018; bootstrap test), but category decoding accuracy during the entire trial or after the response time threshold was not (“all” and “> RT”; p = 0.192 and 0.234). Error bars depict the 95% confidence interval of the mean.

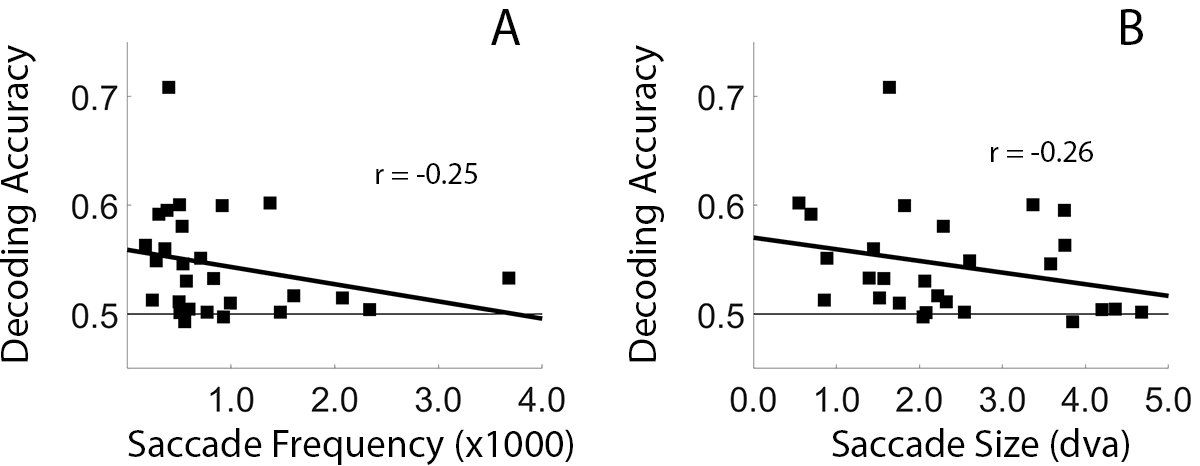

**Figure S3. Individual Differences in Decoding Accuracy Plotted as a Function of Saccade Frequency (A) and Saccade Size (B) during Experiment 1.** Neither correlation was statistically significant.

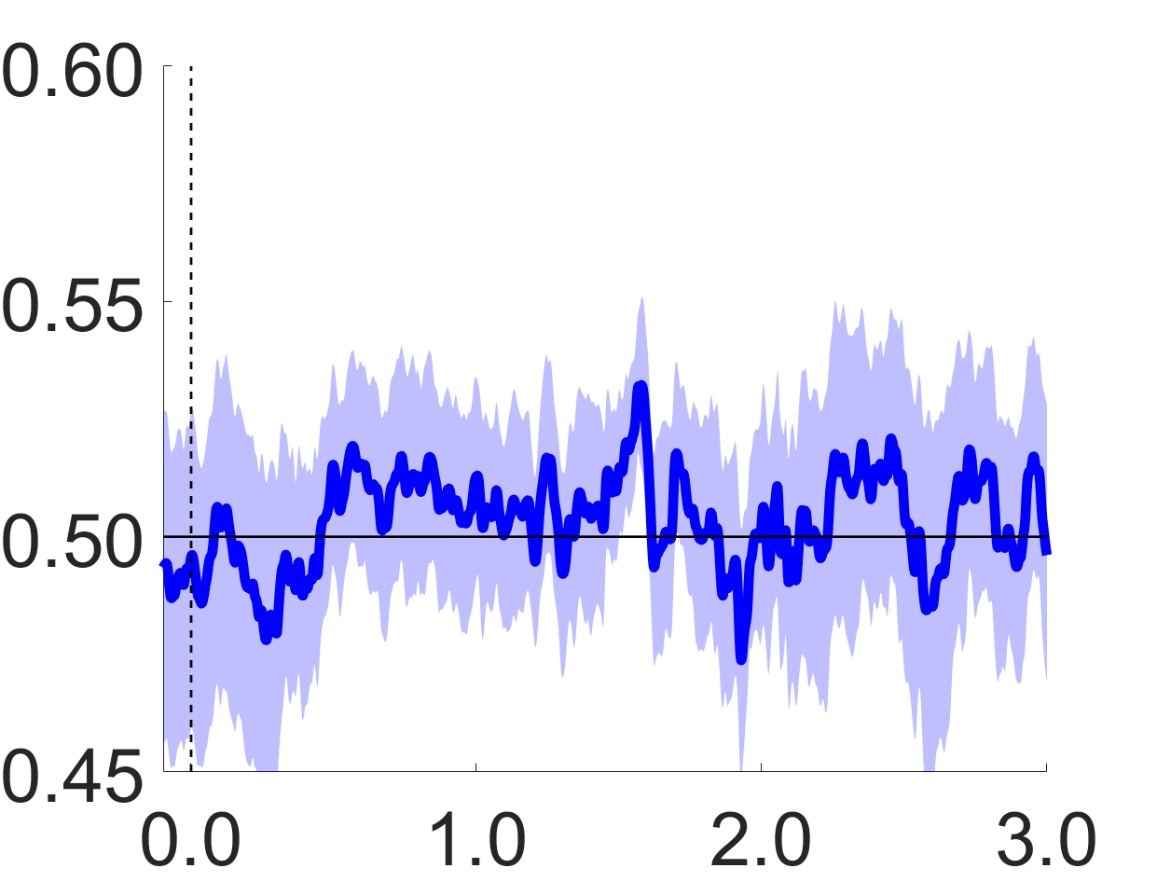

**Figure S4. Response decoding performance as a function of time during Experiment 1**. We trained a linear classifier to predict category membership from unambiguous trials (i.e., ±2°, ±5°, ±15°, ±45°) and used the trained classifier to decode participants responses (i.e., Category 1 vs. Category 2) during ambiguous trials (±0°). This analysis failed to reveal above-chance decoding accuracy at any point during the 3.0 second trial. Shaded regions depict the 95% confidence interval of the mean.
